# E3 ubiquitin ligase SKP2 limits autophagy during *S. aureus* infection

**DOI:** 10.1101/2025.11.03.686196

**Authors:** Abhishek K Singh, Madina Baglanova, Eylin Topfstedt, Kristin Surmann, Silva Holtfreter, Michael Lammers, Barbara M. Bröker, Karsten Becker, Clemens Cammann, Ulrike Seifert

**Affiliations:** Friedrich Loeffler-Institute of Medical Microbiology, University Medicine Greifswald, Greifswald, Germany; Department of Functional Genomics, Interfaculty Institute for Genetics and Functional Genomics, University Medicine Greifswald, Greifswald, Germany; Institute of Immunology, University Medicine Greifswald, Greifswald, Germany; Department of Synthetic and Structural Biochemistry, Institute of Biochemistry, University of Greifswald, Greifswald, Germany

## Abstract

Ubiquitination is a posttranslational modification that affects protein function, stability, and localization and is thereby balancing protein homeostasis. During infection, ubiquitination is crucial in regulating host cell signaling pathways in pathogen recognition, clearance and mounting an efficient immune response. *S. aureus* is an opportunistic pathogen that is able to invade and multiply within both phagocytic and non-phagocytic mammalian cells depending on virulence factor expression of the respective *S. aureus* strain. Selective autophagy serves as a host defense mechanism to combat intracellular bacterial persistence by targeting and degrading intracellular pathogens. However, *S. aureus* can subvert autophagosomal degradation and exploit these organelles for intracellular replication.

We examined the role of the E3 ligase S-phase kinase-associated protein 2 (SKP2), a component of the SKP1-Cullin1-F-box (SCF) – complex, during *S. aureus* infection in alveolar epithelial and in macrophage-like cells. Upon *S. aureus* infection, we demonstrate increased SKP2 abundance through acetylation-induced stabilization and translocation into the cytoplasm. Cytoplasmic SKP2 modulated autophagy induction. By downregulation of SKP2, the level of the autophagy marker LC3-II was elevated which was accompanied by increased survival of intracellular *S. aureus*. Conversely, SKP2 overexpression in host cells reduced LC3-II levels followed by reduced intracellular bacteria. These findings underscore that SKP2 is an important regulator of autophagy, preventing excessive autophagy from being exploited by *S. aureus*. In conclusion, our findings reveal novel molecular mechanisms involved in the interaction between host cells and *S. aureus* providing potential approaches for targeted therapeutic intervention.

**Importance:** *Staphylococcus aureus* is a major pathogen responsible for a wide range of human and animal infections and toxin-mediated syndromes. For many years, *S. aureus* was mainly recognized as an extracellular pathogen. However, it has been shown that *S. aureus* can survive in epithelial cells and phagocytes. Intracellular persistence complicates the treatment by protecting the bacteria from antibiotics and immune detection. Elimination of pathogens from intracellular compartments relies on cellular mechanisms such as recognition and ubiquitination by host E3 ligases followed by selective autophagy. *S. aureus* has developed strategies to manipulate this process by evading degradation by interfering with autophagosome formation or even exploiting the formation of autophagosomes as replication niches. The present study reveals an intricate cellular response pathway that is involved in the intracellular recognition and elimination of the pathogen. This knowledge opens new avenues towards targeted treatment.

## Introduction

*Staphylococcus aureus* is a Gram-positive opportunistic pathogen that causes a wide range of diseases from minor skin infections to life-threatening diseases including sepsis, pneumonia, endocarditis, toxic shock syndrome, and osteomyelitis (1). While *S. aureus* frequently colonizes human epithelial and mucosal surfaces without causing symptoms, it possesses the remarkable ability to invade and replicate within both phagocytic and non-phagocytic mammalian cells (2–4). This intracellular lifestyle contributes to its persistence and complicates treatment efforts (5). *S. aureus* is one of the leading bacteria-related cause of deaths worldwide. By 2021, *S. aureus*-related lower respiratory infections caused 424,000 deaths, representing a 67.6% increase since 1990 (6). These numbers highlight the urgent need to address *S. aureus* as a critical and escalating public health concern.

The ubiquitin-proteasome system (UPS) is an essential component of the host’s defense arsenal against pathogens such as *S. aureus*. During pathogen invasion, host cellular signaling networks experience significant ubiquitin-dependent modifications which, include the activation of innate immune responses, the restructuring of cellular organelles, the reorganization of the cytoskeleton, and metabolic reprogramming aimed at limiting proliferation of the pathogen and/or its elimination (7, 8). The UPS also plays a critical role in the cellular host defense by tagging intracellular bacteria for selective autophagy, thereby influencing the intracellular fate of *S. aureus* (9, 10). During ubiquitination, E3 ligases, which confer substrate specificity in the process of ubiquitination, regulate the fate and function of proteins by attaching chains of ubiquitin moieties to substrate proteins via specific lysine residues (11, 12). Several E3 ligases have been shown to regulate autophagy by influencing autophagosome formation, cargo recruitment, and degradation pathways (7). We recently demonstrated that the E3 ligase LRSAM1 is required for the ubiquitination of *S. aureus* and the subsequent targeting of the bacteria to selective autophagy (13). Ubiquitinated bacteria are recognized by autophagy receptors such as p62/SQSTM1 and NDP52 binding both ubiquitinated bacteria and microtubule-associated protein light chain 3 (LC3), facilitating the delivery of *S. aureus* to autophagosomes for degradation (14). Thus, autophagy serves as a conserved lysosomal degradation pathway and a host defense mechanism. *S. aureus,* however, subverts this process by exploiting autophagosomes as replication niche (15, 16). Further, *S. aureus* manipulates autophagosome dynamics to evade degradation and facilitate its intracellular persistence (17, 18).

S-phase kinase-associated protein 2 (SKP2), a member of the F-box protein family, is a critical component of the SKP1-Cullin1-F-box (SCF) complex, which targets various substrates for ubiquitin-mediated proteasomal degradation (19, 20). SKP2 is well known for its role in cell cycle regulation through ubiquitination and subsequent proteasomal degradation of the cyclin-dependent kinase inhibitors p27, p21 and p57 (20–23). Through its E3 ligase activity, SKP2 controls the stability of multiple proteins involved in key cellular pathways both via K48 and K68 linked ubiquitination (23). SKP2 expression is tightly regulated under normal conditions; however, in disease states most notably in cancer, it can be transcriptionally upregulated (24–28). Additionally, post-translational modifications like acetylation and phosphorylation of SKP2 have been described, which affect its activity, stability, and subcellular localization (29–33). Moreover, it has been shown that SKP2 plays a second, context-dependent role in autophagy, acting either as a suppressor or as a promoter. The E3 ligase can inhibit autophagy by marking key regulators like PHLPP1 and Beclin1 for degradation. This contributes to tumor cell survival, therapy resistance, as well as to disease progression in viral infection and fibrosis (34–36).

While the role of SKP2 in cancer regulation is well documented, its function in the regulation of autophagy in response to intracellular bacteria has not been addressed so far. This study aims to investigate the role of SKP2 in the host cellular response during *S. aureus* infection, thereby uncovering novel autophagic functions of SKP2 that promote host defense and pathogen clearance.

## Results

### Increased abundance of SKP2 upon *S. aureus* infection in human lung epithelial cells

To investigate the role of SKP2 in the host cell response to *S. aureus*, we first examined whether SKP2 expression is modulated upon bacterial infection. We selected A549 cells, a human alveolar epithelial cell line that is commonly used as an *in vitro* model for respiratory infections, because of their relevance in studying host-pathogen interactions including intracellular pathogens (37, 38). For infection, we used *S. aureus* USA300 a major methicillin-resistant *Staphylococcus aureus* (MRSA) clone of high clinical relevance, which is known for its ability to invade or persist within host cells (39, 40). To eliminate extracellular bacteria while preserving those that had successfully invaded the host cells, lysostaphin was applied after one hour of infection.

Immunoblot analysis of the infected cells showed an increase of SKP2 levels 2 h, 3 h and 5 h post infection (Figure 1A and B). Infection also led to rapid degradation of IκBα within the first hours, with protein levels gradually restored over time, indicating that the NF-κB pathway was activated (Figure 1A and C). Cytokine determination in cell culture supernatants of infected cells showed increased concentrations of IL-6, significant at 5 h post infection, while IL-8 secretion increased progressively throughout the infection time course (Figure 1D and E). Lactate dehydrogenase (LDH) was also elevated, confirming infection-induced cytotoxicity (Figure 1F).

**Figure 1.**
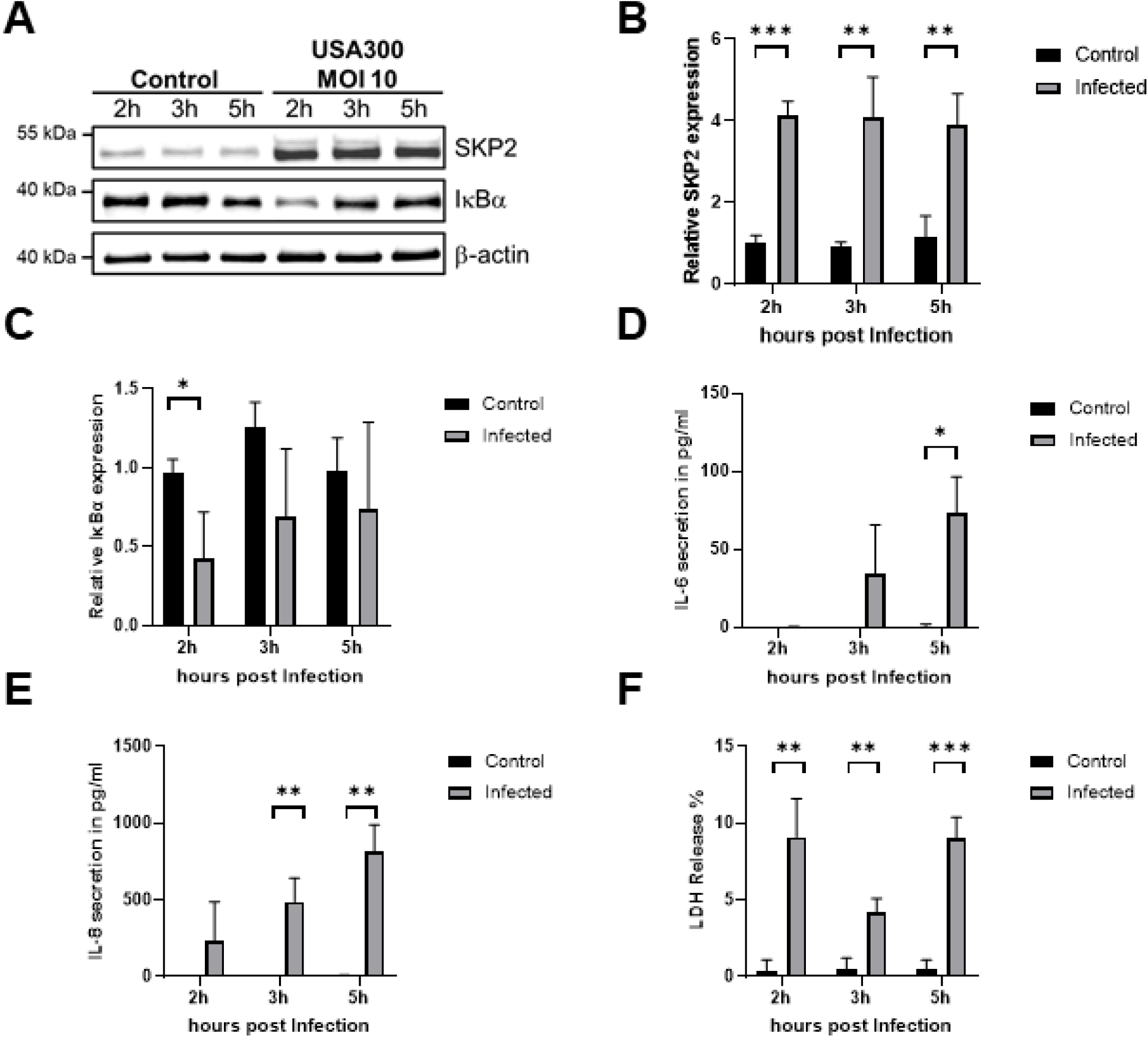
Increased abundance of SKP2 upon *S. aureus* infection in A549 cells. A549 cells were infected with *S. aureus* USA300 (MOI 10) for the depicted time points. SKP2 and IκBα abundance was analyzed by immunoblot compared to uninfected controls (A). Densitometric analysis of SKP2 and IκBα normalized to β-actin and relative expression was calculated to the 2 h uninfected control, n = 3 (B, C). Cytokine secretion was assessed via ELISA for IL-6, n = 3 (D) and IL-8, n = 3 (E). Cytotoxicity was monitored by determining extracellular LDH at 2 h, 3 h, and 5 h post-infection, n = 3 (F). Data in B-F are presented as mean ± SD (*p < 0.05; **p < 0.01; ***p < 0.001, students t-test).

To confirm that the accumulation of SKP2 was not specific to A549 cells, we also infected primary small airway epithelial cells (SAECs), which represent naive airway epithelium and macrophage-like THP-1 cells, which serve as model for professional phagocytes involved in innate immune responses. A similar accumulation of SKP2 2 h, 3 h and 5 h post infection was observed in SAEC cells (Supplementary Figure 1A and B) and THP-1 cells (Supplementary Figure 2A and B). However, there were subtle differences between the cell lines in their reaction to *S. aureus* infection. NF-κB activation as reflected by reduced IκBα levels was prolonged in THP-1 cells when compared to A549 cells and SAEC (Supplementary Figures 1A and C and 2A and C). SAECs had high basal IL-6 secretion and secreted more IL-6 upon infection than A549 cells. They were also more resistant to cell death. However, high basal cytokine secretion in non-infected SAECs may reflect a cell type–specific effect, potentially influenced by the growth conditions under which the cells were cultivated (Supplementary Figure 1D - F). In THP-1 cells a strong IL-1β release indicated pyroptosis induction, which was also reflected in higher cell death shown by increased LDH release (Supplementary Figure 2D and E). These results indicate that the increase in SKP2 is a general process observed in different cell types upon infection with *S. aureus*.

### Cellular localization of SKP2 upon *S. aureus* infection

Subsequent analysis of SKP2 expression in the context of *S. aureus* infection revealed a disconnection between SKP2 mRNA and protein levels. While SKP2 protein levels were elevated in infected epithelial cells, the corresponding mRNA levels decreased slightly (Figure 2A). These results indicated that the SKP2 protein might be stabilized by post-translational modifications in response to *S. aureus* infection.

**Figure 2.**
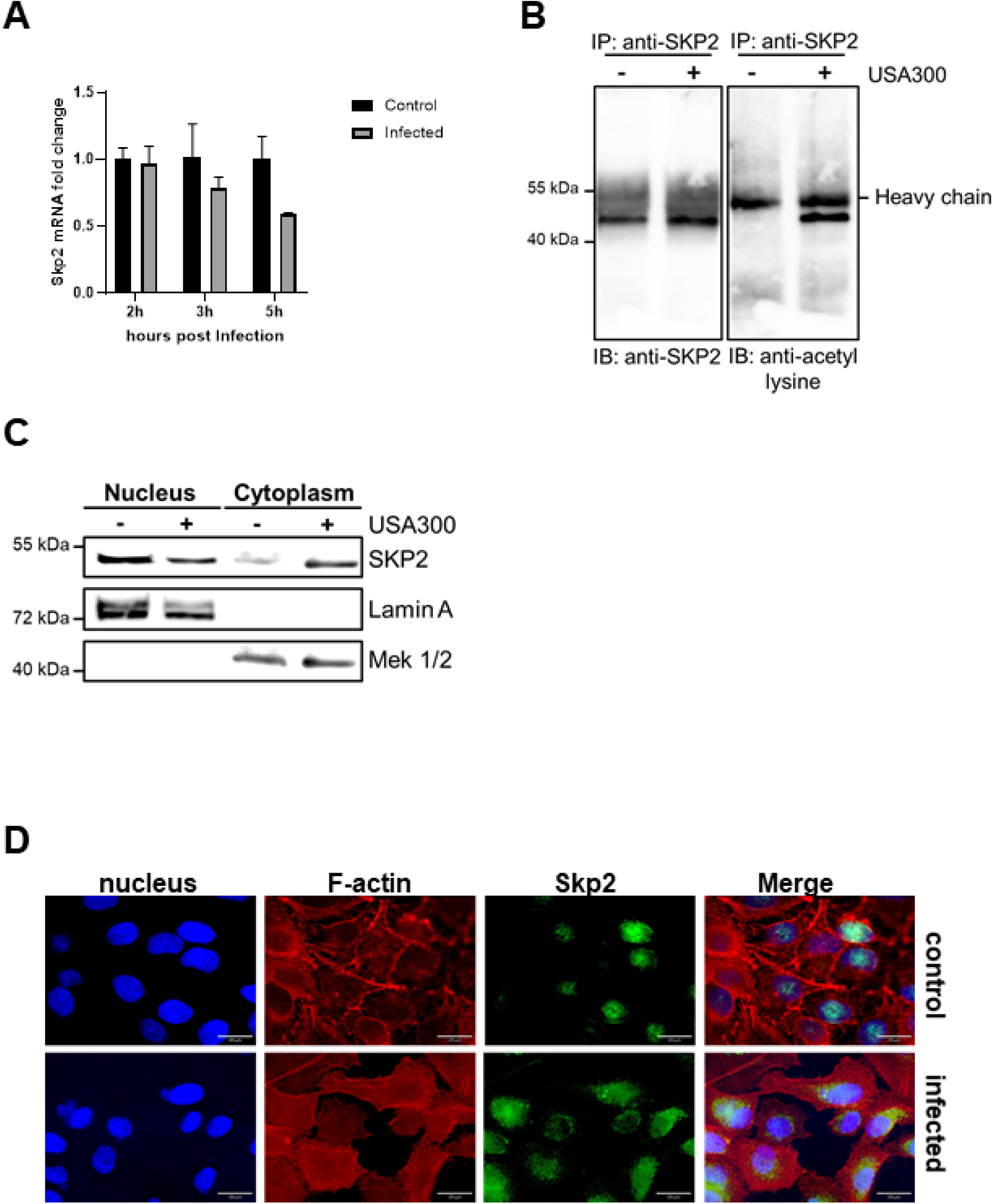
Regulation and localization of SKP2 upon *S. aureus* infection. Changes in *SKP2* mRNA were analyzed by qPCR in A549 cells infected with *S. aureus* USA300 (MOI 10) at the depicted time points, fold change was calculated to 2 h uninfected control and normalized to the *RPL47* housekeeping gene, n = 3 (A). SKP2 was immunoprecipitated from A549 cells infected with *S. aureus* USA300 (MOI 10) for 5 h and analyzed with anti-SKP2 and anti-acetyl lysine antibody compared to uninfected controls, heavy chain is indicated as unspecific band from the precipitating SKP2 antibody, n = 2 (B). Analysis of nuclear and cytoplasmic fractionation at 5 h post infection with *S. aureus* USA300 (MOI 10) with Lamin A serving as control for the nuclear fraction and MEK 1/2 serving as control for the cytoplasmic fraction, n = 2 (C). Immunofluorescence analysis of A549 cells infected for 5 h with *S. aureus* USA300 (MOI 10). Nuclei are stained with DAPI (blue), F-actin filaments with phalloidin (red), and SKP2 with an anti-SKP2-Alexa488 antibody (green), magnification 60x, scalebar 20 µm, n = 2 (D).

Because acetylation of SKP2 prevents its proteasomal degradation in cancer cells (29–33), we examined whether a similar effect occurs in response to *S. aureus* infection. Following immunoprecipitation of SKP2 from infected A549 and THP-1 cells, we detected lysine acetylation by immunoblotting (Figure 2B and Supplementary Figure 3A). This suggests that acetylation-mediated stabilization of SKP2 contributes to its increased abundance during infection.

Next, we asked whether *S. aureus* infection affects the subcellular localization of SKP2. Under non-infected conditions, SKP2 exhibited dual subcellular localization in the cytoplasm and nucleus, with the majority located in the nucleus (41). However, upon infection-induced acetylation, a translocation of SKP2 from the nucleus to the cytoplasm was observed, as evidenced by both subcellular fractionation with immunoblotting and immunofluorescence microscopy (Figures 2C and D). These data suggest that the *S. aureus* infection triggers a shift of SKP2 from its nuclear localization to the cytoplasm, implicating a potential alteration in its cellular function in response to *S. aureus*.

### *S. aureus* protein A (SpA) contributes to increased SKP2

Next, we addressed the question what triggers SKP2 stabilization during *S. aureus* infection. To this end, A549 cells were treated with different concentrations of recombinant *S. aureus* protein A (SpA), a multi-functional virulence factor expressed by the pathogen (42).

Treatment with SpA from 1 h to 6 h revealed a rapid increase in SKP2 abundance similar to that observed upon infection with *S. aureus* suggesting a role of SpA in modulating SKP2 protein levels (Figure 3A and B). To confirm this, we infected the cells with the USA300Δ*spa* strain that is deficient in SpA and compared SKP2 levels to cells infected with the wild-type USA300 strain. As expected, the wild-type strain induced a significantly higher abundance of SKP2 in host cells after 5 h infection than USA300Δ*spa* (Figure 3C and D).

**Figure 3.**
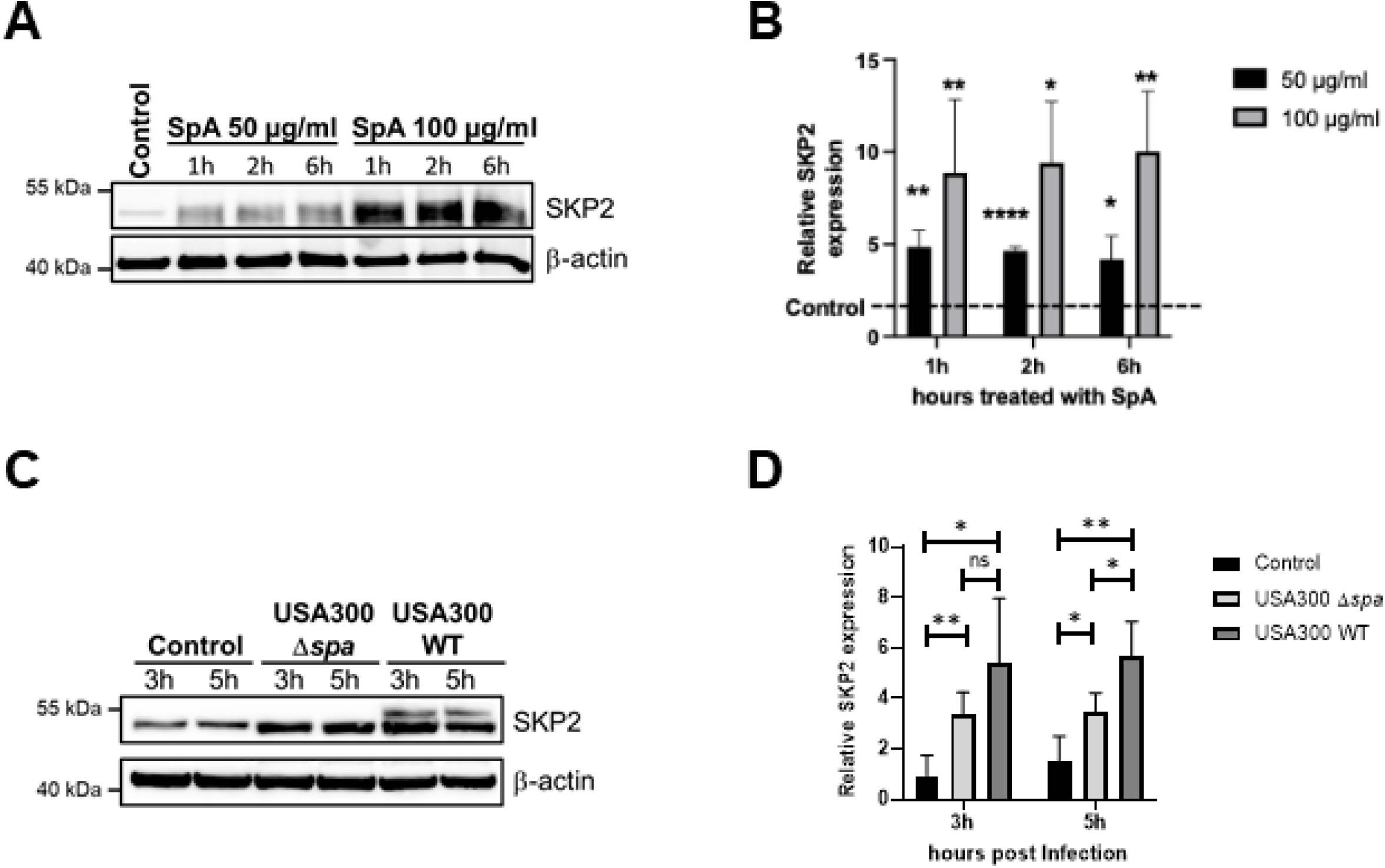
*S. aureus* protein A (SpA) contributes to increased SKP2 levels. A549 epithelial cells treated with 50 µg / ml and 100 µg / ml SpA for the depicted timepoints were analyzed for SKP2 levels by immunoblotting with β-actin as loading control (A). Densitometric analysis of SKP2 from immunoblot data, normalized to β-actin. Relative expression was calculated to the untreated control, which is indicated by the dotted line, n = 3 (B). A549 epithelial cells were infected with *S. aureus* USA300 (MOI 10) and USA300Δ*spa* (MOI 10) for the depicted time points and analyzed for SKP2 expression by immunoblotting with β-actin as loading control (C). Densitometric analysis of SKP2 expression normalized to β-actin, relative expression was calculated to the 3 h uninfected control, n = 4 (D). Data in B and D are presented as mean ± SD (*p < 0.05; **p < 0.01; ****p < 0.0001, students t-test).

### SKP2 regulates intracellular *S. aureus* survival and autophagy

To investigate the role of SKP2 in regulating cellular processes, siRNA-mediated knockdown of SKP2 was performed in A549 cells. The high virulence of *S. aureus* USA300 complicated the analysis because the bacteria induced substantial host cell death upon SKP2 knockdown. Therefore, we utilized the *S. aureus* strain HG001, which was less cytotoxic, enabling a thorough investigation of the effects of SKP2 knockdown on bacterial survival. There was a substantial increase in intracellular bacterial colony-forming units (CFUs) upon SKP2 knockdown compared to the infected non-targeting control (NTC) cells 5 h post infection (Figure 4A). Despite the stabilization of residual SKP2 in the siRNA-treated cells following infection, a substantial accumulation of the SKP2 substrate protein, cell cycle inhibitor p27, could be observed, indicating a substantial decrease in SKP2 activity within these cells (Figure 4B - D).

**Figure 4.**
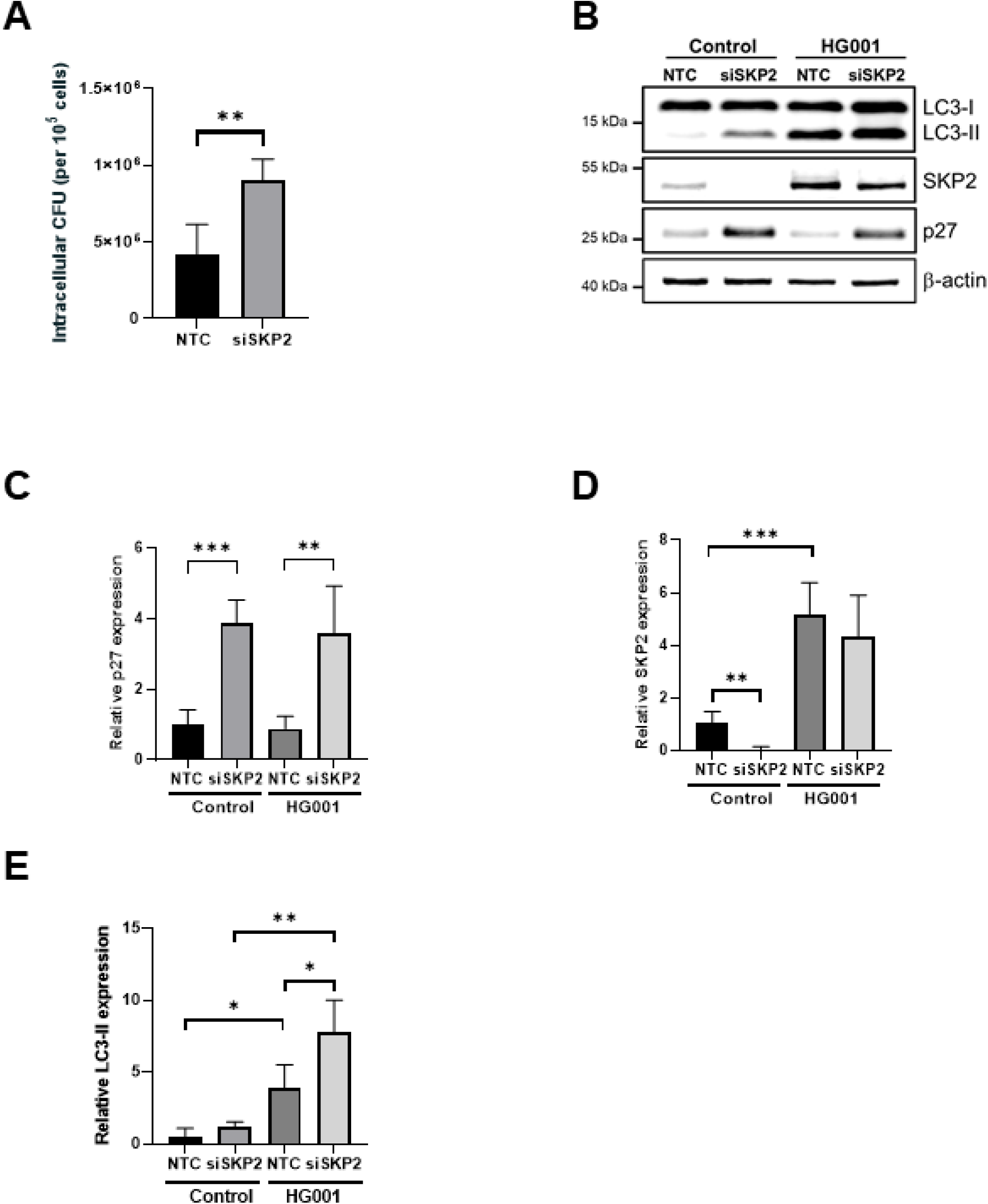
Increased intracellular *S. aureus* and autophagy induction upon siRNA-mediated reduction of SKP2. A549 cells treated with SKP2-siRNA (siSKP2) or non-targeted control (NTC) siRNA were infected with *S. aureus* HG001 (MOI 20) for 5 h. Intracellular bacterial burden was determined as CFU normalized to 1 × 10⁵ host cells, n = 4 (A). Cell lysates were analyzed for LC3-I, LC3-II, SKP2, and the SKP2 target p27 as a knockdown control, by immunoblotting with β-actin as loading control (B). Densitometric analysis of p27, SKP2, and LC3-II normalized to β-actin, relative expression was calculated to NTC control, n = 4 (C - E). Data in A, C-E are presented as mean ± SD (*p<0.05, **p<0.01, ***p<0.001 students t-test).

A similar increase in intracellular CFUs could be obtained by treating the cells with the SKP2-specific inhibitor SZL-P141 (Supplementary Figure 3B). Moreover, co-application of the inhibitor to siRNA-treated cells further amplified the intracellular CFU counts (Supplementary Figure 3B). However, the increase in the number of intracellular live *S. aureus* had no effect on secretion of the cytokines IL-6 and IL-8 by the infected host cells (Supplementary Figure 3C and D).

To explain the observed increase in intracellular bacteria upon SKP2 knockdown, we aimed to ascertain the precise function of SKP2 during the infection process. SKP2 plays an important role during cell cycle progression. A previous study suggested that *S. aureus* invasion slows down host cell proliferation due to a specific delay in the G2/M phase transition, which benefits intracellular bacterial survival (43). We therefore investigated if a similar cell cycle delay upon SKP2 inhibition was causing a higher intracellular bacterial load. However, single or combined SKP2-knockdown and inhibition had no effect on cell cycle progression in *S. aureus*-infected cells (Supplementary Figure 3E).

Given the translocation of SKP2 to the cytoplasm following infection, we next investigated the impact of SKP2 knockdown on selective autophagy. The protein LC3 serves as a key marker of autophagy, with LC3-II specifically indicating the formation of autophagosomes (44). Upon siRNA-mediated downregulation of SKP2, we observed an increase in LC3-II formation, indicating an upregulation of autophagy (Figure 4B and E). This suggests that SKP2 may interfere with autophagy during infection.

To further confirm this, we generated an expression vector with SKP2 carrying the K145R and K228R point mutations (pcDNA3.1 SKP2^K145R^ ^K228R^) to prevent degradation by proteasomes. The mutated amino acid residues correspond to ubiquitination sites of SKP2 according to mUbiSiDa, a comprehensive database for mammalian protein ubiquitination sites (45). Overexpression of SKP2 in A549 cells did not result in a robust increase on protein level (data not shown). However, we observed a consistent higher abundance of SKP2^K145R^ ^K228R^ compared to SKP2 wildtype and transfected controls in HeLa cells (Supplementary Figure 4A). Therefore, we had to focus our subsequent experiments on HeLa cells, to investigate the impact of SKP2 overexpression on autophagy and intracellular *S. aureus* survival. As *S. aureus* is known to elicit cell-type-specific responses we first tested whether HeLa cells exhibit a comparable increase in SKP2 levels upon infection. To this end, we infected HeLa cells with *S. aureus* HG001 (MOI 10 and 20) for 2 h, 3 h and 5 h and observed a similar increase in SKP2 to that seen in A549 cells (Supplementary Figure 4B and C). To determine whether SKP2 overexpression modulates autophagy during bacterial infection, HeLa cells were transfected with the pcDNA3.1 SKP2^K145R^ ^K228R^ vector and subsequently infected with *S. aureus* HG001. The number of viable *S. aureus* was significantly reduced in cells overexpressing SKP2 ^K145R^ ^K228R^ in comparison with the controls, confirming SKP2’s contribution to the elimination of intracellular bacteria (Figure 5A). SKP2 overexpression also markedly reduced LC3-II accumulation upon infection, confirming that high levels of SKP2 suppress *S. aureus*-induced autophagy (Figure 5B - D). To further investigate whether high levels of SKP2 inhibit autophagy under non-infectious conditions as well, we assessed the formation of LC3-II upon treatment with rapamycin alone or in combination with bafilomycin A1 (46). Rapamycin, an mTOR inhibitor, induces autophagy by promoting autophagosome formation, whereas bafilomycin A1 blocks autophagosome-lysosome fusion, preventing autophagic degradation and leading to autophagosome accumulation. Rapamycin treatment alone increased LC3-II lipidation in HeLa cells, indicating enhanced autophagosome formation, which was even increased upon addition of bafilomycin indicating a high autophagic flux (Figure 5E and F). Although the overexpression of SKP2^K145R^ ^K228R^ had no effect on basal levels of LC3-II, the induction of autophagy using rapamycin resulted in a reduction in LC3-II lipidation in these cells compared to the control cells. When treated with a combination of rapamycin and bafilomycin the transfection with the SKP2^K145R^ ^K228R^ mutant reduced the levels of LC3-II compared with transfection with the empty vector. This clearly shows that SKP2 downregulates autophagy (Figure 5E and F).

**Figure 5.**
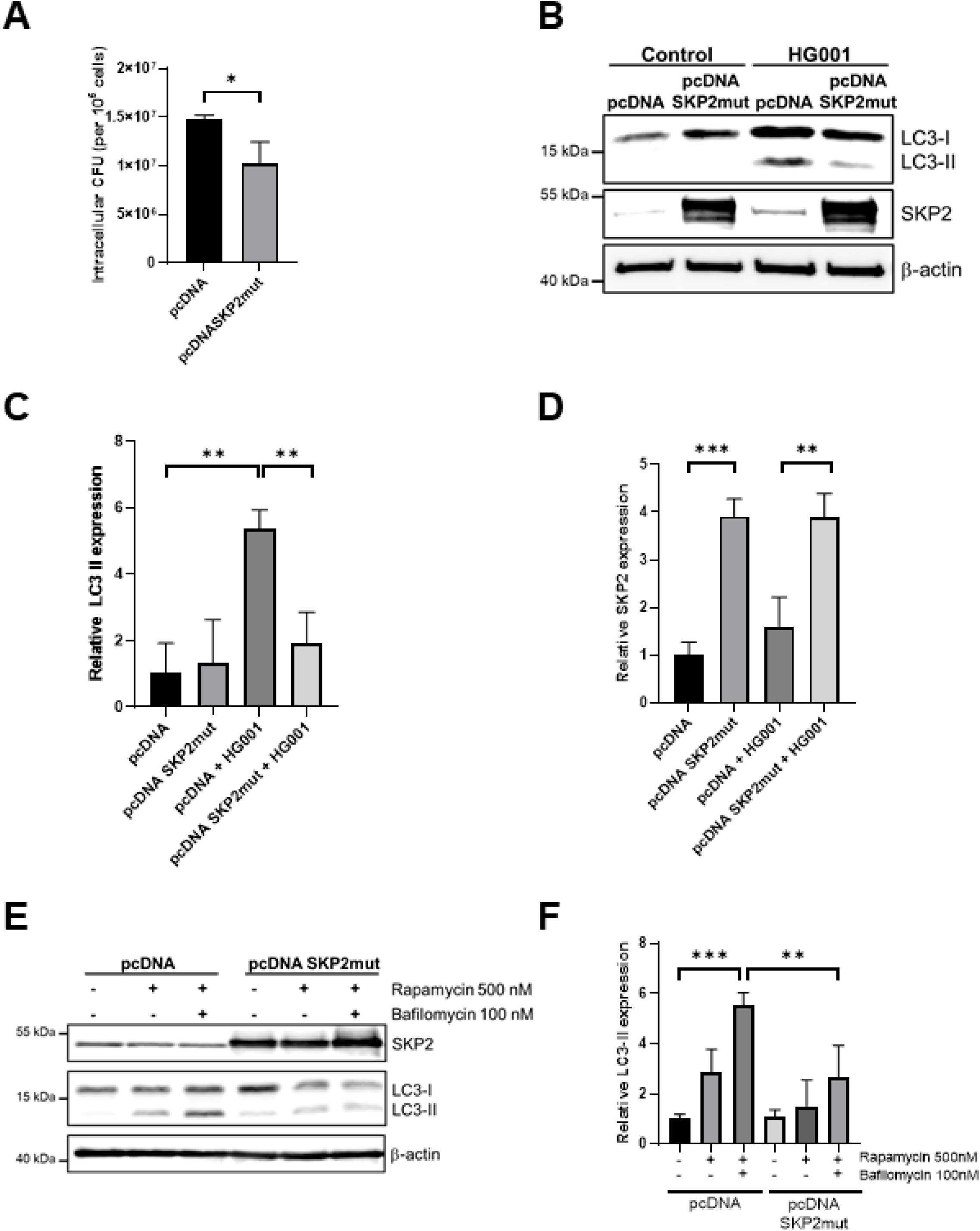
Reduced intracellular *S. aureus* and formation of LC3-II in HeLa cells expressing SKP2^K145R^ ^K228R^. HeLa cells transfected for 24 h with SKP2^K145R^ ^K228R^ (pcDNA SKP2mut) or empty vector control (pcDNA) were infected with *S. aureus* HG001 (MOI 20) for 3 h. Intracellular bacterial burden was determined as CFU normalized to 1 × 10⁵ host cells, n = 3 (A). SKP2, LC3-I and LC3-II abundance was assessed via immunoblot with β-actin as loading control (B). Densitometric analysis of LC3-II and SKP2 normalized to β-actin, relative expression was calculated to the uninfected control, n = 3 (C, D). HeLa cells transfected for 24 h with SKP2^K145R^ ^K228R^ (pcDNA SKP2mut) or empty vector control (pcDNA) were treated with 500 nM rapamycin alone or in combination with 100 nM bafilomycin A1 for 6 h or left untreated and subsequently analyzed for SKP2, LC3-I and LC3-II abundance via immunoblot with β-actin as loading control (E). Densitometric analysis of LC3-II formation normalized to β-actin, relative expression was calculated to the untreated vector control, n = 3 (F). Data are presented as mean ± SD. Data in A, C, D and F are presented as mean ± SD (*p<0.05, **p<0.01, ***p<0.001 students t-test).

In summary, these results indicate a prominent role of SKP2 in regulating autophagy, enabling the elimination of intracellular *S. aureus* while preventing excessive autophagy to be exploited by *S. aureus*.

## Discussion

Epithelial cells represent the first line of defense against *S. aureus* by initiating immune responses such as cytokine release. Accumulating evidence highlights *S. aureus* ability to adopt an intracellular phase, especially also in non-professional phagocytes such as epithelial cells, endothelial cells, and fibroblasts. Upon internalization *S. aureus* can have two intracellular fates, it can reside within phagosomes or escape into the cytoplasm. This duality in localization reflects a complementary strategy to modulate host responses and to optimize its survival.

In this study, we observed that infection with *S. aureus* led to an increase in SKP2 protein levels in A549 lung epithelial cells, primary small airway epithelial cells (SAECs), HeLa cells, and macrophage-like THP-1 cells. Notably, this upregulation was not driven by increased transcription of SKP2, but rather by post-translational modification through acetylation in A549 and THP1 cells, highlighting the role of acetylation in stabilizing SKP2 expression levels. Histone-acetyltransferase p300 has been shown to acetylate SKP2 lysine residues within nuclear localization signals (NLS) which regulate subcellular distribution of SKP2 in cancers such as prostate cancer cells (47). We demonstrated that this mechanism is also induced by *S. aureus* during infection. The infected cells exhibited a shift in SKP2 localization, which was consistent with acetylation-mediated translocation to the cytoplasm. Consistently, we observed no changes in cell cycle regulation upon SKP2 inhibition in infected A549 cells reflecting the cytoplasmic localization of SKP2 during infection (Supplementary Figure 3E). In addition to regulating its intracellular localization, acetylation of SKP2 stabilizes the protein by interfering with its degradation. SKP2 is normally targeted for its proteasomal degradation by the E-Cadherin (Cdh1)-associated anaphase-promoting complex/cyclosome (APC/C), an E3 ubiquitin ligase complex (33). Importantly, the region of SKP2 encompassing amino acids 46 to 90, containing the acetylated lysine residues K68 and K71, is critical for its interaction with Cdh1 (33). Acetylation at these sites likely disrupts this interaction, thereby impairing Cdh1-mediated ubiquitination and subsequent degradation by proteasomes. This can explain how infection with *S. aureus* increases the total cellular amount of SKP2. The histone-acetyltransferase p300 cooperates with the Yes-associated protein (YAP), which has recently been implicated as a key regulator of both autophagy and inflammation as it activates transcription at the enhancer and promoter regions of relevant target genes (48, 49). It is, therefore, plausible that p300’s acetylation of SKP2 contributes to the modulation of autophagy during *S. aureus* infection, either directly by interfering with SKP2’s Cdh1 binding or indirectly by affecting its subcellular localization.

Furthermore, our data indicates that the observed increase in SKP2 may be caused by *S. aureus* protein A (Figure 3). The surface adhesin SpA is a well-characterized virulence factor known to play multifaceted roles in *S. aureus* pathogenesis, including the promotion of immune suppression (50). Interestingly, infection with the USA300Δ*spa* strain did not result in complete attenuation of SKP2 expression, suggesting that additional virulence factors may contribute to this cellular response such as α-hemolysin, which has been reported to interfere with cellular host responses (51, 52). SpA is known to interact with the epidermal growth factor receptor (EGFR) and the tumor necrosis factor receptor 1 (TNFR1) in epithelial cells and macrophages (53, 54). Activation of these receptors has previously been shown to modulate nuclear p300 levels, e.g. by EGFR-mediated upregulation of p300 expression (55), or TNFR1-mediated activation of the NF-κB pathway, which enhances p300 activity (56, 57). Given the role of p300 in acetylation of SKP2 these pathways could potentially influence SKP2 expression. However, the precise molecular mechanisms linking receptor engagement to SKP2 regulation remain to be elucidated.

As mentioned before, upon internalization by epithelial cells such as A549, *S. aureus* can have two distinct intracellular fates: confinement within phagosomal compartments via selective autophagy or escape into the cytosol. *S. aureus* uses multiple strategies to manipulate the host autophagy pathway. It induces the formation of LC3-positive autophagosomes in a variety of host cells, ranging from professional phagocytes (58, 59) to epithelial cells (60, 61). Remarkably, inhibiting autophagy has been shown to reduce intracellular *S. aureus* survival (62). This process is further reinforced by blocking autophagic flux, thereby promoting bacterial persistence and replication (63). Previous research has shown, that rapamycin-induced autophagy restores both replication and cytotoxicity in *agr*-deficient *S. aureus* strains, indicating that autophagy is crucial for bacterial replication, cytoplasmic escape, and host cell killing (64). Our experiments show that SKP2 overexpression can attenuate rapamycin-induced autophagy as well as autophagy caused by *S. aureus* infection. Moreover, inhibition of autophagy results in reduced bacterial survival (Figure 5 and 64, 16). This capacity to adopt compartment-specific strategies highlights a flexible intracellular program aimed at evading degradation while maintaining access to nutrient-rich environments. These findings suggest that *S. aureus* not only survives autophagic processes but actively utilizes them to maintain intracellular persistence.

We propose that the increase in SKP2 levels in infected cells represents a host cell mechanism to limit autophagy, thereby preventing its exploitation by *S. aureus* for survival and virulence. To investigate this putative role, we performed infections with siRNA-mediated knockdown of SKP2 in A549 cells which resulted in enhanced autophagosome formation, as evidenced by the accumulation of the lipidated form of LC3 (LC3-II). Upon *S. aureus* infection, LC3-II was markedly elevated, especially in siRNA-treated cells, suggesting a strong autophagosome accumulation, which was accompanied by a significant increase in intracellular bacterial load. Consistent with this, pharmacological inhibition of SKP2 with SZL-P141 similarly elevated the numbers of intracellular vital bacteria, which was further enhanced when combining SKP2 knockdown and inhibitor treatment (Supplementary Figure 3B). Conversely, SKP2 overexpression reduced the formation of LC3-II indicating lower levels of autophagy. This was accompanied by less intracellular bacterial survival. These results support the hypothesis that SKP2 acts as negative regulator of autophagy thereby limiting intracellular *S. aureus* survival.

The notion that *S. aureus* can exploit autophagosomes as replicative niches is particularly intriguing. Autophagosomes provide a membranous compartment that shields the bacterium from recognition by the host cells. While nutrient availability within the autophagosome is limited, *S. aureus* has a flexible metabolism and may utilize available lipids or amino acids to sustain growth. Despite the higher intracellular bacterial load, the siRNA-treated cells showed no increased secretion of the proinflammatory cytokines IL-6 and IL-8. It is plausible that within autophagosomes the bacteria evade the detection by host cytosolic pattern recognition receptors (PRRs) (65). Epithelial cells such as A549 possess a limited PRR repertoire, largely restricted to the cell surface. It has been speculated that this reflects an evolutionary trade-off, because epithelia prioritize tissue integrity over robust immune defense as found in professional immune cells (66).

Considering that SKP2 inhibition facilitates intracellular persistence of *S. aureus*, pharmacological inhibition of SKP2 may inadvertently compromise host antimicrobial defense mechanisms. This is particularly critical in immunocompromised individuals and cancer patients, where innate immunity is often impaired. Notably, bortezomib, an FDA-approved proteasome inhibitor widely used in the treatment of multiple myeloma also functions as a SKP2 inhibitor, although its approval was based on broader proteasome-inhibiting properties (67, 68). Its therapeutic success has sparked growing interest in developing more selective SKP2 inhibitors with reduced off-target effects. However, our findings underscore a potential risk: targeting SKP2 may increase the susceptibility to intracellular bacterial infections in vulnerable individuals. As more specific SKP2-targeting therapies are developed, it will be essential to evaluate their impact on host immunity and infection susceptibility. Conversely, and in line with this, a recent study also highlighted SKP2’s protective role in sepsis-induced acute lung injury. Intravenous administration of SKP2 mRNA-encapsulating lipid nanoparticles significantly improved lung function and reduced mortality in septic mice, underscoring the therapeutic potential of SKP2 (69).

In summary, our observations reveal a nuanced host-pathogen interplay wherein the upregulation of SKP2 by the host cells prevents excessive induction of autophagy, thereby – counterintuitively – limiting intracellular *S. aureus* survival. Obviously, *S. aureus* can subvert host autophagy pathways for its own benefit. A deeper understanding of these mechanisms may pave the way for innovative therapies that target the host-pathogen interface and extend the anti-bacterial therapeutic portfolio to meet an urgent clinical need.

## Materials and Methods

### Cell lines

The human alveolar epithelial cell line A549 (70) and the cervical cancer cell line HeLa (ATCC #CCL-2) were cultured in RPMI 1640 medium (Gibco) supplemented with 10% fetal calf serum (FCS; Capricorn). Small Airway Epithelial Cells (SAEC; Lonza) were cultured according to the manufacturer’s instructions using SAGM™ BulletKit™ medium (Lonza). THP-1 monocytes (ATCC #TIB-202) were cultured in RPMI 1640 medium (Gibco) with 10% FCS (Capricorn), stimulated with 100 ng/mL phorbol 12-myristate 13-acetate (PMA) (Sigma-Aldrich) for 24 h to induce differentiation, and subsequently rested for 48 h prior to further use.

Staphylococcal protein A (SpA, Sigma), rapamycin (Sigma), bafilomycin A1 (Selleckchem) and SZL-P141 (Selleckchem) were used for cell treatment throughout the study at the indicated concentrations and time points described below.

### *S. aureus* propagation

*S. aureus* USA300, derived from strain JE2, is a community-associated methicillin-resistant *S. aureus* (CA-MRSA) strain (71). *S. aureus* USA300Δ*spa*, a kind gift of Jan Marten van Dijl, is a transposon mutant of JE2, with a deletion of the *spa* gene, which encodes protein A (72).*S. aureus* HG001 strain derived from the widely studied *S. aureus* strain NCTC8325 (73).All strains were maintained in tryptic soy broth (TSB, Roth) 37 °C and 220 rpm or on blood agar plates (BD Biosciences) at 37°C. For cellular internalization assays, *S. aureus* was initially grown in TSB as a pre-culture for 2 h until reaching an optical density (OD_600_) of 0.8. Subsequently, the bacteria were inoculated and cultured into RPMI medium supplemented with 10% fetal calf serum (FCS) or SAGM™ BulletKit™ medium and grown until an OD_600_ of 0.35 was achieved.

### *S. aureus* internalization assay

The infection protocol was adapted from a protocol described before to meet the criteria of the present study (13, 74, 75). Cells were infected with *S. aureus* at a multiplicity of infection (MOI) of 10. For *S. aureus* internalization, an infection master mix was prepared by combining the respective strain of *S. aureus* (in RPMI with 10% FCS) with 2.9 μl of 7.5% NaHCO_3_ per ml of bacterial culture. This mixture was added to the cells, with a determined cell number and the co-incubation was carried out at 37 °C with 5% CO_2_ for 1 h to allow bacterial internalization. During this incubation, 100 μl of the infection master mix were diluted in PBS without Mg²⁺ or Ca²⁺ and plated on blood agar to determine the bacterial count, which was later used to calculate the precise MOI achieved. Following the 1-hour incubation, the infection master mix was removed, and pre-warmed RPMI containing lysostaphin (Amibion) (final concentration: 10 μg / ml) was added. Cell cultures were then incubated again at 37 °C and 5% CO₂. At different time points post lysostaphin treatment the culture medium was removed, and the cells were washed with PBS. Cells were collected in 500 µl of TRIzol™ reagent (Thermo Fisher Scientific) for subsequent immunoblotting.

#### CFU determination

Cells were lysed at indicated time-points post infection through hypotonic lysis using bidistilled water (*Aqua bidest.*) for 30 min. Serial dilutions from 10^-3^ – 10^-5^ of the released bacteria in *Aqua bidest.* were plated on blood agar plates. CFU counts were determined after 24 h incubation at 37°C and the mean CFU numbers were calculated from all dilutions. For normalization, a technical replicate of the infected cells was harvested using 1% trypsin (Capricorn Scientific) and subsequently counted using trypan blue staining to exclude dead cells in a counting chamber (Neubauer). Finally, CFUs were normalized to 1 x 10^5^ viable cells.

#### Immunoblot analysis

Proteins were extracted using TRIzol™ reagent following the manufacturer’s guidelines and quantified via Bradford assay. For immunoblotting, proteins were resolved by SDS-PAGE, transferred onto nitrocellulose membranes, and probed with primary antibodies against SKP2, IκBα, LC3, p27, β-actin, lamin A, MEK1/2 (all from Cell Signaling Technology), anti-acetyl lysine antibody (Abcam) and subsequently detected using species-specific HRP conjugated secondary antibodies (Dianova). For the densitometric analysis of SKP2, both bands were considered, referring to the possible existence of SKP2 isoforms (28). Membranes were visualized using chemiluminescence with SignalFire™ ECL Reagent (Cell Signaling Technology) and analyzed with the ImageQuant 800 system (Cytiva).

#### Immunoprecipitation

Cell lysates (100–200 µg total protein) were prepared using radioimmunoprecipitation assay (RIPA) lysis buffer supplemented with protease inhibitors. For immunoprecipitation, 1 µL (1 µg) of SKP2 antibody (Invitrogen) was added to each lysate and incubated overnight at 4°C on a rotator. Magnetic beads (100 µL of 10 mg/mL Sure Beads Protein G, BioRad) were equilibrated and washed three times with PBS containing 0.1% Tween, then incubated with the antibody-lysate mixture for 4 h at 4°C. After incubation, beads were separated using a magnetic rack and subsequently washed three times with lysis buffer to remove unbound material. For sample preparation, beads were resuspended in SDS sample buffer, heated at 95°C for 5 min and centrifuged at 14,000–16,000 g for 1 min. The prepared samples were further analyzed via immunoblotting.

#### Cell viability assay

Cell viability was assessed by measuring the LDH content in cell supernatants using the CytoTox-ONE Homogeneous Membrane Integrity Assay (Promega) following the manufacturer’s instructions. Briefly, 50 µL of supernatant were incubated with 50 µL of substrate solution for 10 min. The reaction was then halted by adding 25 µL of Stop solution. Supernatants from non-infected cells treated for 10 minutes at 37°C with 10% TritonX 100 (AppliChem) were used as a positive control to represent 100% cell death. Samples were read at 590 nm using a plate reader (Tecan).

#### Cytokine analysis

ELISA for IL-6, IL-8 and IL-1β (all BioLegend) was performed according to manufacturer’s instructions. In brief, plates were coated with the included capture antibody and incubated for 16-18 h at 4°C. After washing (3 times, PBS + 0.5% Tween-20). Plates were blocked for 1 h at 500 rpm, washed, and incubated with standards or samples for 2 h with shaking. After washing, plates were treated with biotinylated detection antibody (1 h), avidin-HRP (30 min), and TMB substrate (15 min, dark), followed by stop solution (2 N H₂SO₄) and absorbance measurement at 450 nm.

#### Generation of SKP2^K145R^ ^K228R^ mutant

For overexpression experiments the SKP2 construct was cloned into the mammalian overexpression vector pcDNA3.1Myc/His (Thermo Fisher Scientific) using the pOTB7-SKP2 vector provided by the mammalian gene collection (Dharmacon) as template. Mutations for lysine to arginine at position 145 and 228 were introduced using the Q5® Site-Directed Mutagenesis Kit (New England Biolabs) according to the manufacturer’s description using the following mutagenesis primers K145R_fwd 5’-CCTCACAGGTCGCAATCTGCACC – 3’ and K145R_rev 5’-TCTAAGGTCTGCCATAG - 3’; K228R_fwd 5’-TACTCTCGCACGCAACTCAAATTTAGTGC -3’ and K228R_rev 5’-TTGACAATGGGATCCG - 3’. The obtained pcDNA3.1 SKP2^K145R^ ^K228R^ constructs were transformed into chemically competent *E. coli* and verified by Sanger sequencing (Eurofins Genomics).

#### Transfection of pcDNA3.1 SKP2^K145R^ ^K228R^ and SKP2-siRNA

Cells were seeded 24 h prior to transfection at a density of 2 × 10⁵ cells per well in 6-well plates. The following day, cells were transfected with either plasmid DNA for overexpression pcDNA3.1 myc/his SKP2 (pcDNA SKP2), pcDNA3.1 myc/his SKP2^K145R^ ^K228R^ (pcDNA SKP2mut) or empty vector control pcDNA3.1 myc/his; 5 µg per well or with siRNA for knockdown (SKP2-siRNA or non-targeting control siRNA, Dharmacon) using ScreenFect® A (ScreenFect GmbH) according to the manufacturer’s instructions. DNA/siRNA and ScreenFect A were diluted in ScreenFect buffer, mixed, and incubated for 20 min at room temperature to allow complex formation. Complexes were then added dropwise to cells cultured in RPMI medium without serum. After 4 h, the medium was supplemented with RPMI containing 10% FCS, and cells were incubated under standard conditions. HeLa cells were infected 24 h post-transfection, whereas A549 cells were allowed to rest for 48 h before infection.

#### Statistical Analysis

Densitometric analysis of the immunoblot results was performed using ImageJ software (version 1.54g; National Institutes of Health, USA). Experiments were presented as means +/- SD and Student’s t-test was used to compare differences between analyzed samples. All statistical analyses were performed using GraphPad Prism software, version 9, and were considered significant at \**P* < 0.05, ** *P* < 0.01, *** *P* < 0.001, and **** *P* < 0.0001.

#### Funding

Funded by the Deutsche Forschungsgemeinschaft (DFG, German Research Foundation) – project number(s); RTG 2719)

